## Supplementary material for "Network-principled deep generative models for designing drug combinations as graph sets": SI

#### 1 Methods

##### 1.1 More details on HVGAE – Embedding disease-disease network

While we briefly discussed the details of the HVGAE for disease-disease network in section 3.1.2 of the paper, here we provide a more detailed formulation. Given the adjacency matrix $A^{(\mathrm{d})}$, and the node attributes for diseas-disease network, which is derived by attentional pooling and denoted by $\hat{F}^{(\mathrm{d})}$, the encoder for this level of HVGAE is defined as:

$$
\begin{aligned}
q(\boldsymbol{Z}^{(\mathrm{d})} \,|\, \hat{F}^{(\mathrm{d})}, A^{(\mathrm{d})}) = \prod_{i=1}^{n_\mathrm{d}} q(\boldsymbol{z}_i^{(\mathrm{d})} \,|\, \hat{F}^{(\mathrm{d})}, A^{(\mathrm{d})}), \quad \text{where} \quad q(\boldsymbol{z}_i^{(\mathrm{d})} \,|\, \hat{F}^{(\mathrm{d})}, A^{(\mathrm{d})}) = \mathcal{N}(\boldsymbol{\mu}_i^{(\mathrm{d})}, diag(\boldsymbol{\sigma}_i^{2,(d)})),\\
\boldsymbol{\mu}^{(\mathrm{d})} = \mathrm{GNN}_{\boldsymbol{\mu},d}(A^{(\mathrm{d})}, \hat{F}^{(\mathrm{d})}) \quad \in \mathbb{R}^{n_\mathrm{d} \times L_\mathrm{d}},\\
\log(\boldsymbol{\sigma}^{(\mathrm{d})}) = \mathrm{GNN}_{\boldsymbol{\sigma},d}(A^{(\mathrm{d})}, \hat{F}^{(\mathrm{d})}) \quad \in \mathbb{R}^{n_\mathrm{d} \times L_\mathrm{d}}.
\end{aligned} \tag{1}
$$

In the equation above, $\mathrm{GNN}_{\boldsymbol{\mu},d}$ and $\mathrm{GNN}_{\boldsymbol{\sigma},d}$ are multi-layer graph neural networks; $n_\mathrm{d}$ is the number of nodes; $L_\mathrm{d}$ is the dimension of latent variables; and $\boldsymbol{\mu}^{(\mathrm{d})}$ and $\boldsymbol{\sigma}^{(\mathrm{d})}$ are matrices of mean vectors and standard deviation vectors, respectively.
The generative model is given by inner product decoder between latent variable. More specifically,

$$
p(A^{(\mathrm{d})} \,|\, \boldsymbol{Z}^{(\mathrm{d})}) = \prod_{i=1}^{n} \prod_{j=1}^{n} p(A_{ij}^{(\mathrm{d})} \,|\, \boldsymbol{z}_i^{(\mathrm{d})}, \boldsymbol{z}_j^{(\mathrm{d})}), \quad \text{where} \quad p(A_{ij}^{(\mathrm{d})} \,|\, \boldsymbol{z}_i^{(\mathrm{d})}, \boldsymbol{z}_j^{(\mathrm{d})}) = \sigma(\boldsymbol{z}_i^{(\mathrm{d})} \boldsymbol{z}_j^{(\mathrm{d})T}), \tag{2}
$$

with $\sigma(\cdot)$ as the logistic sigmoid function. Finally, the evidence lower bound (ELBO) for this level of HVGAE is defined as follows:

$$
\mathcal{L}^{(\mathrm{d})} = \mathbb{E}_{q(\boldsymbol{Z}^{(\mathrm{d})} \,|\, \hat{F}^{(\mathrm{d})}, A^{(\mathrm{d})})}[\log p(A^{(\mathrm{d})} \,|\, \boldsymbol{Z}^{(\mathrm{d})})] - \mathrm{KL}\big(q(\boldsymbol{Z}^{(\mathrm{d})} \,|\, \hat{F}^{(\mathrm{d})}, A^{(\mathrm{d})}) \,||\, p(\boldsymbol{Z}^{(\mathrm{d})})\big), \tag{3}
$$

##### 1.2 More details on RL – Policy Network

In the main text Sec. 3.2.3, we have described the policy network adopted for the reinforcement learning agent. Here we detail how a link prediction-based action is made. Specifically, each component is sampled according to a predicted distribution governed by the following equations (?):

$$a_t^{(k)} = \text{CONCAT}(a_{\text{fr},t}^{(k)}, a_{\text{sc},t}^{(k)}, a_{\text{et},t}^{(k)}, a_{\text{tr},t}^{(k)}),$$
$$a_t = \{a_t^{(k)}\}_{k=1}^{K},$$
$$X_t = \{X_t^{(k)}\}_{k=1}^{K},$$

where

$$\begin{aligned} f_1(s_t) &= \text{SOFTMAX}(\text{FC}_{\text{fr}}(X_t, Y)), \\ a_{\text{fr,t}}^{(k)} &\sim f_1(s_t) \in \{0,1\}^{n_k}; \\ f_2(s_t) &= \text{SOFTMAX}(\text{FC}_{\text{sc}}(X_{a_{\text{fr},t}}, X_t, Y)), \\ a_{\text{sc,t}}^{(k)} &\sim f_2(s_t) \in \{0,1\}^{(n_k+c)}; \\ f_3(s_t) &= \text{SOFTMAX}(\text{FC}_{\text{et}}(X_{a_{\text{fr},t}}, X_{a_{\text{sc},t}}, Y)), \\ a_{\text{et,t}}^{(k)} &\sim f_3(s_t) \in \{0,1\}^{b}; \\ f_4(s_t) &= \text{SOFTMAX}(\text{FC}_{\text{tr}}(X_t, Y)), \\ a_{\text{tr,t}}^{(k)} &\sim f_4(s_t) \in \{0,1\}; \end{aligned} \tag{4}$$

where FC's are fully connected neural networks, subscripts 'fr' and 'sc' indicate the first and the second drug, and $c$ is the cardinality of $C$. Also, $Y$ is the targeted disease we are generating drug combination for.

#### 1.3 Drug-drug interactions considered

Table S1: Various drug-drug interactions considered in this study, with individual probabilities predicted (Ryu *et al.*, 2018) for designed pairs. Positive or negative annotations were manually made.

| | | |
|---|---|---|
| 1 | Drug a can cause a decrease in the absorption of Drug b resulting in a reduced serum concentration and potentially a decrease in efficacy. | N |
| 2 | Drug a can cause an increase in the absorption of Drug b resulting in an increased serum concentration and potentially a worsening of adverse effects. | N |
| 3 | The absorption of Drug b can be decreased when combined with Drug a. | N |
| 4 | The bioavailability of Drug b can be decreased when combined with Drug a. | N |
| 5 | The bioavailability of Drug b can be increased when combined with Drug a. | P |
| 6 | The metabolism of Drug b can be decreased when combined with Drug a. | N |
| 7 | The metabolism of Drug b can be increased when combined with Drug a. | P |
| 8 | The protein binding of Drug b can be decreased when combined with Drug a. | N |
| 9 | The serum concentration of Drug b can be decreased when it is combined with Drug a. | N |
| 10 | The serum concentration of Drug b can be increased when it is combined with Drug a. | P |
| 11 | The serum concentration of the active metabolites of Drug b can be increased when Drug b is used in combination with Drug a. | P |
| 12 | The serum concentration of the active metabolites of Drug b can be reduced when Drug b is used in combination with Drug a resulting in a loss in efficacy. | N |
| 13 | The therapeutic efficacy of Drug b can be decreased when used in combination with Drug a. | N |
| 14 | The therapeutic efficacy of Drug b can be increased when used in combination with Drug a. | P |
| 15 | Drug a may decrease the excretion rate of Drug b which could result in a higher serum level. | P |
| 16 | Drug a may increase the excretion rate of Drug b which could result in a lower serum level and potentially a reduction in efficacy. | N |
| 17 | Drug a may decrease the cardiotoxic activities of Drug b. | P |
| 18 | Drug a may increase the cardiotoxic activities of Drug b. | N |
| 19 | Drug a may increase the central neurotoxic activities of Drug b. | N |
| 20 | Drug a may increase the hepatotoxic activities of Drug b. | N |

| | | |
|---|---|---|
| 21 | Drug a may increase the nephrotoxic activities of Drug b. | N |
| 22 | Drug a may increase the neurotoxic activities of Drug b. | N |
| 23 | Drug a may increase the ototoxic activities of Drug b. | N |
| 24 | Drug a may decrease effectiveness of Drug b as a diagnostic agent. | N |
| 25 | The risk of a hypersensitivity reaction to Drug b is increased when it is combined with Drug a. | N |
| 26 | The risk or severity of adverse effects can be increased when Drug a is combined with Drug b. | N |
| 27 | The risk or severity of bleeding can be increased when Drug a is combined with Drug b. | N |
| 28 | The risk or severity of heart failure can be increased when Drug b is combined with Drug a. | N |
| 29 | The risk or severity of hyperkalemia can be increased when Drug a is combined with Drug b. | N |
| 30 | The risk or severity of hypertension can be increased when Drug b is combined with Drug a. | N |
| 31 | The risk or severity of hypotension can be increased when Drug a is combined with Drug b. | N |
| 32 | The risk or severity of QTc prolongation can be increased when Drug a is combined with Drug b. | N |
| 33 | Drug a may decrease the analgesic activities of Drug b. | N |
| 34 | Drug a may decrease the anticoagulant activities of Drug b. | N |
| 35 | Drug a may decrease the antihypertensive activities of Drug b. | N |
| 36 | Drug a may decrease the antiplatelet activities of Drug b. | N |
| 37 | Drug a may decrease the bronchodilatory activities of Drug b. | N |
| 38 | Drug a may decrease the diuretic activities of Drug b. | N |
| 39 | Drug a may decrease the neuromuscular blocking activities of Drug b. | N |
| 40 | Drug a may decrease the sedative activities of Drug b. | P |
| 41 | Drug a may decrease the stimulatory activities of Drug b. | N |
| 42 | Drug a may decrease the vasoconstricting activities of Drug b. | N |
| 43 | Drug a may increase the adverse neuromuscular activities of Drug b. | N |
| 44 | Drug a may increase the analgesic activities of Drug b. | P |
| 45 | Drug a may increase the anticholinergic activities of Drug b. | P |
| 46 | Drug a may increase the anticoagulant activities of Drug b. | P |
| 47 | Drug a may increase the antihypertensive activities of Drug b. | P |
| 48 | Drug a may increase the antiplatelet activities of Drug b. | P |
| 49 | Drug a may increase the antipsychotic activities of Drug b. | P |
| 50 | Drug a may increase the arrhythmogenic activities of Drug b. | N |
| 51 | Drug a may increase the atrioventricular blocking (AV block) activities of Drug b. | N |
| 52 | Drug a may increase the bradycardic activities of Drug b. | N |
| 53 | Drug a may increase the bronchoconstrictory activities of Drug b. | N |
| 54 | Drug a may increase the central nervous system depressant (CNS depressant) activities of Drug b. | N |
| 55 | Drug a may increase the central nervous system depressant (CNS depressant) and hypertensive activities of Drug b. | N |
| 56 | Drug a may increase the constipating activities of Drug b. | N |
| 57 | Drug a may increase the dermatologic adverse activities of Drug b. | N |
| 58 | Drug a may increase the fluid retaining activities of Drug b. | N |
| 59 | Drug a may increase the hypercalcemic activities of Drug b. | N |
| 60 | Drug a may increase the hyperglycemic activities of Drug b. | N |
| 61 | Drug a may increase the hyperkalemic activities of Drug b. | N |
| 62 | Drug a may increase the hypertensive activities of Drug b. | N |
| 63 | Drug a may increase the hypocalcemic activities of Drug b. | N |
| 64 | Drug a may increase the hypoglycemic activities of Drug b. | N |

| | | |
|---|---|---|
| 65 | Drug a may increase the hypokalemic activities of Drug b. | N |
| 66 | Drug a may increase the hyponatremic activities of Drug b. | N |
| 67 | Drug a may increase the hypotensive activities of Drug b. | N |
| 68 | Drug a may increase the hypotensive and central nervous system depressant (CNS depressant) activities of Drug b. | N |
| 69 | Drug a may increase the immunosuppressive activities of Drug b. | N |
| 70 | Drug a may increase the myelosuppressive activities of Drug b. | N |
| 71 | Drug a may increase the myopathic rhabdomyolysis activities of Drug b. | N |
| 72 | Drug a may increase the neuroexcitatory activities of Drug b. | N |
| 73 | Drug a may increase the neuromuscular blocking activities of Drug b. | N |
| 74 | Drug a may increase the orthostatic hypotensive activities of Drug b. | N |
| 75 | Drug a may increase the photosensitizing activities of Drug b. | P |
| 76 | Drug a may increase the QTc-prolonging activities of Drug b. | N |
| 77 | Drug a may increase the respiratory depressant activities of Drug b. | N |
| 78 | Drug a may increase the sedative activities of Drug b. | N |
| 79 | Drug a may increase the serotonergic activities of Drug b. | N |
| 80 | Drug a may increase the stimulatory activities of Drug b. | P |
| 81 | Drug a may increase the tachycardic activities of Drug b. | N |
| 82 | Drug a may increase the thrombogenic activities of Drug b. | N |
| 83 | Drug a may increase the ulcerogenic activities of Drug b. | N |
| 84 | Drug a may increase the vasoconstricting activities of Drug b. | N |
| 85 | Drug a may increase the vasodilatory activities of Drug b. | N |
| 86 | Drug a may increase the vasopressor activities of Drug b. | N |

### 2 Network score results

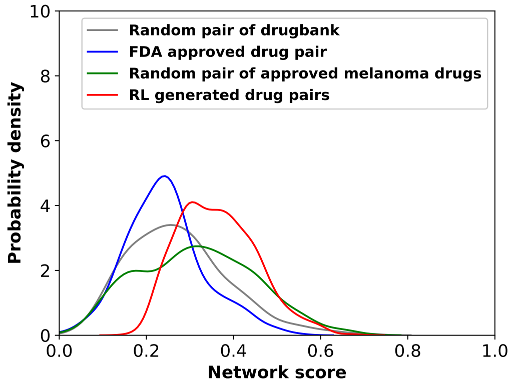


(a) Melanoma

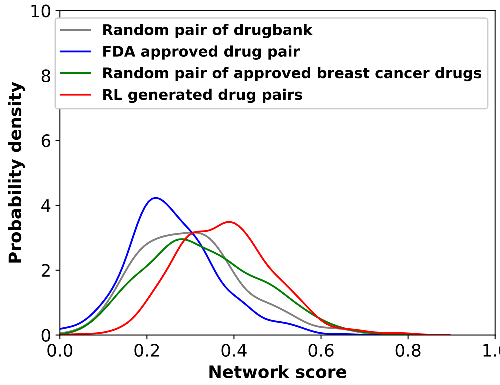


(b) Breast cancer

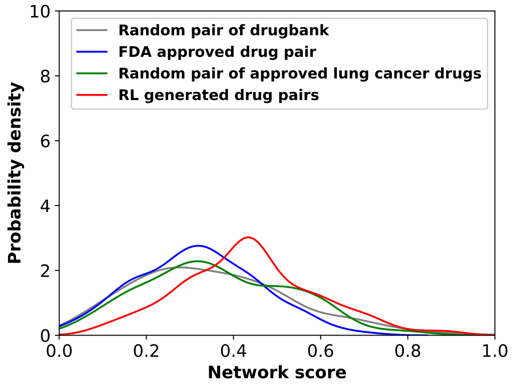


(c) Lung cancer

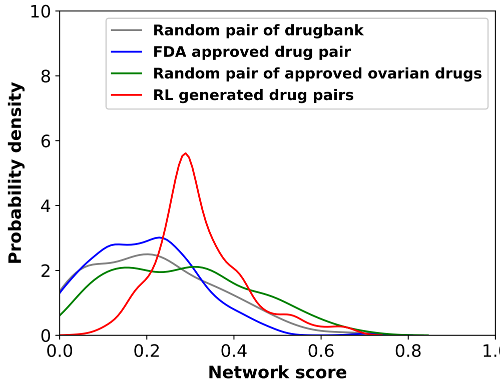


(d) Ovarian cancer

Figure S1: Comparison of distribution of network score between RL generated drug combinations, FDA approved drug combinations, randomly selected drug pairs from drugbank and randomly selected drug pairs from FDA approved single drug for melanoma, lung, breast and ovarian cancers.

Table S2: One-sided KS test statistics for comparison of network score distributions (p-values)

| RL vs. | drugbank | FDA drug comb. | cancer spec. |
|---|---|---|---|
| Melanoma | 1.55 e-37 | 6.05 e-74 | 1.46 e-15 |
| Breast cancer | 7.05 e-7 | 6.47 e-59 | 1.02 e-17 |
| Lung cancer | 1.48 e-27 | 2.22 e-43 | 6.48 e-24 |
| Ovarian cancer | 5.29 e-68 | 1.73 e-90 | 8.28 e-30 |

Table S3: Comparison of percentage of low and high network score

| | Low network score ($< 0.2$) | | | | High network score ($> 0.5$) | | | |
|---|---|---|---|---|---|---|---|---|
| | drugbank | drug pair | Cancer | RL | drugbank | drug pair | Cancer | RL |
| Melanoma | 28% | 31.6% | 23.8% | 0.2% | 4.6% | 0.4% | 7.8% | 6.2% |
| Breast cancer | 18% | 22.6% | 15% | 2.2% | 6% | 2% | 9.4% | 11.6% |
| Lung cancer | 20.6% | 21.6% | 17.6% | 6.4% | 15.4% | 8% | 19.4% | 24.6% |
| Ovarian cancer | 50.2% | 54.2% | 37.6% | 11.2% | 3.6% | 0.4% | 10.6% | 7.2% |

### 3 Toxicity results

#### 3.1 Melanoma

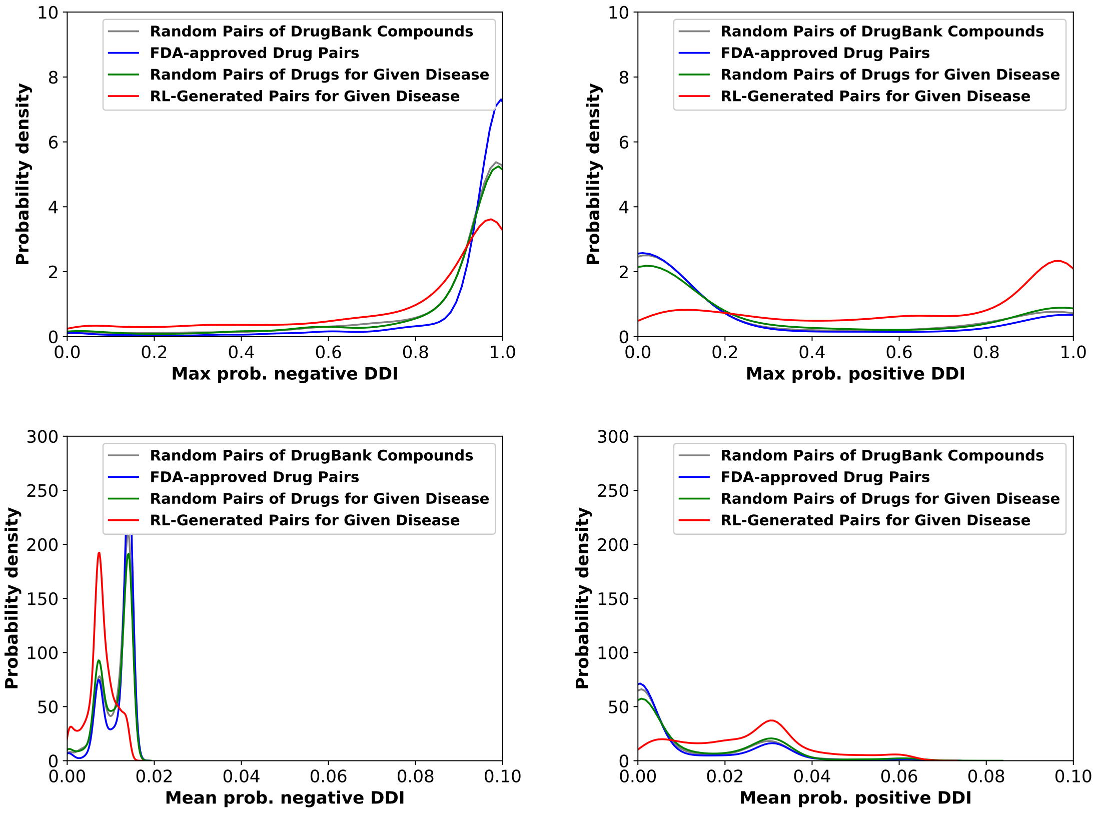


Figure S2: Comparison of distribution of toxicity between RL generated drug combinations, FDA approved drug combinations, randomly selected drug pairs from drugbank and randomly selected drug pairs from FDA approved single drug for melanoma.

#### 3.2 Breast cancer

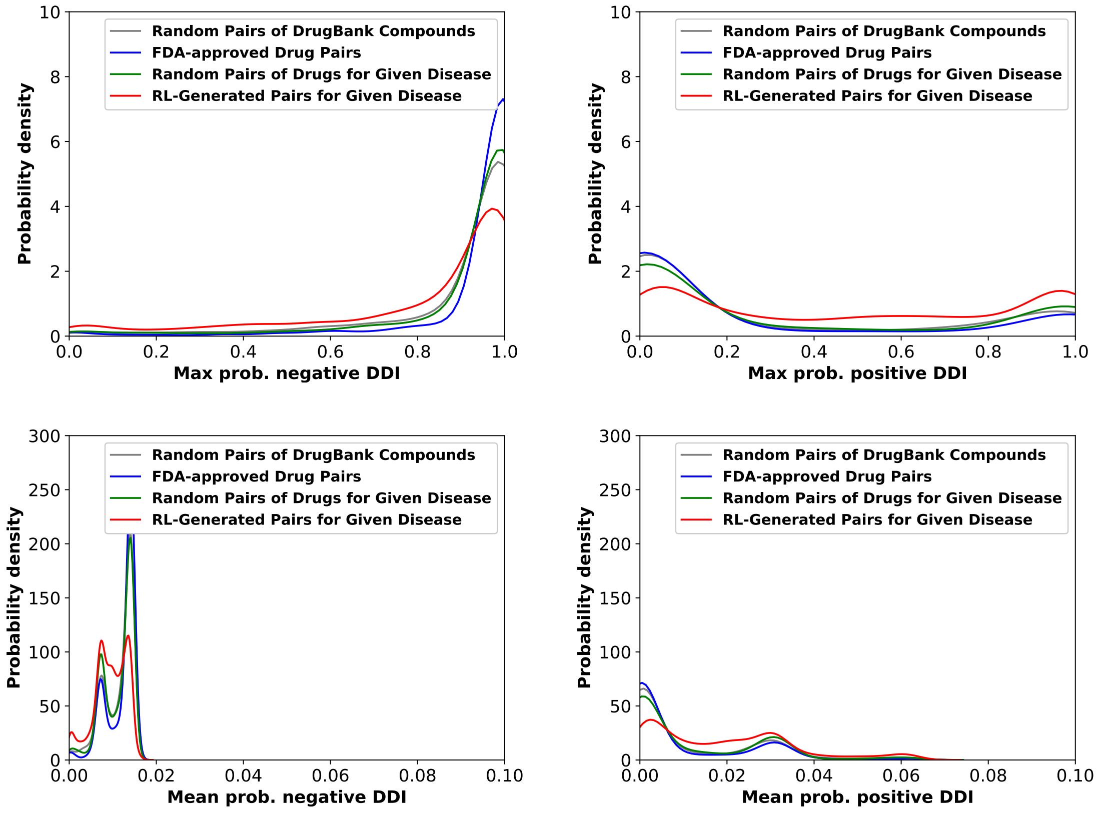


Figure S3: Comparison of distribution of toxicity between RL generated drug combinations, FDA approved drug combinations, randomly selected drug pairs from drugbank and randomly selected drug pairs from FDA approved single drug for breast cancer.

#### 3.3 Lung cancer

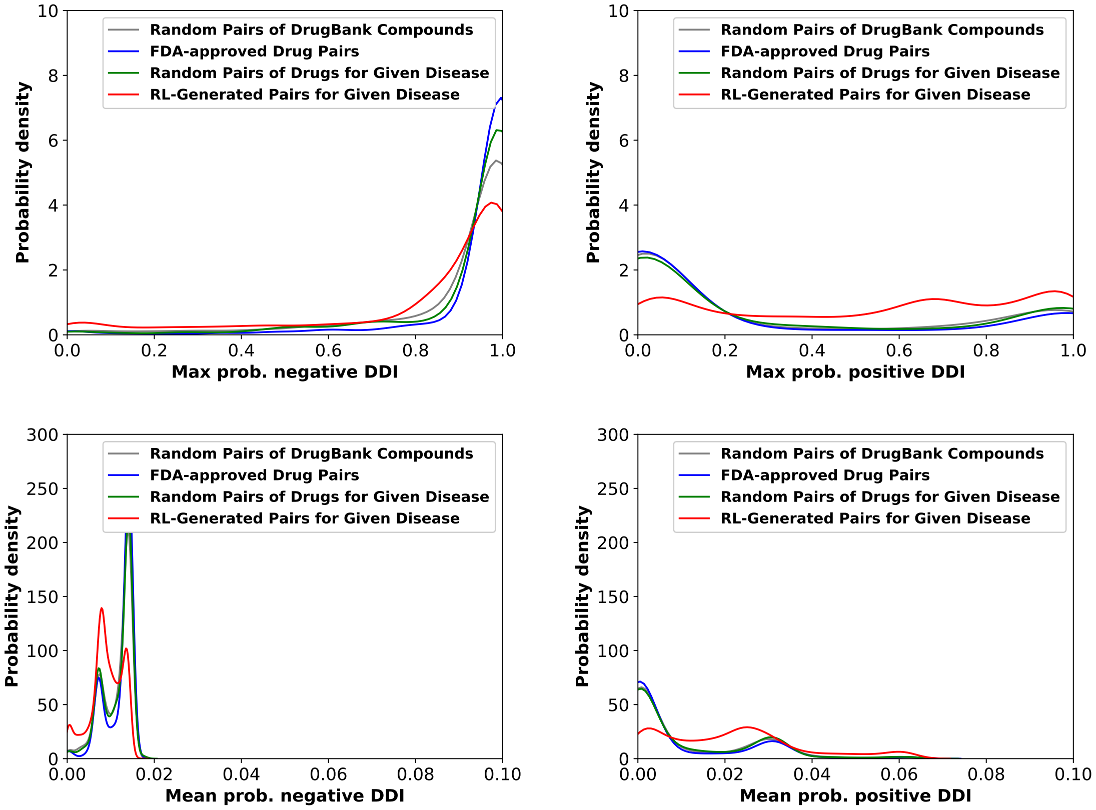


Figure S4: Comparison of distribution of toxicity between RL generated drug combinations, FDA approved drug combinations, randomly selected drug pairs from drugbank and randomly selected drug pairs from FDA approved single drug for lung cancer.

##### 3.4 Ovarian cancer

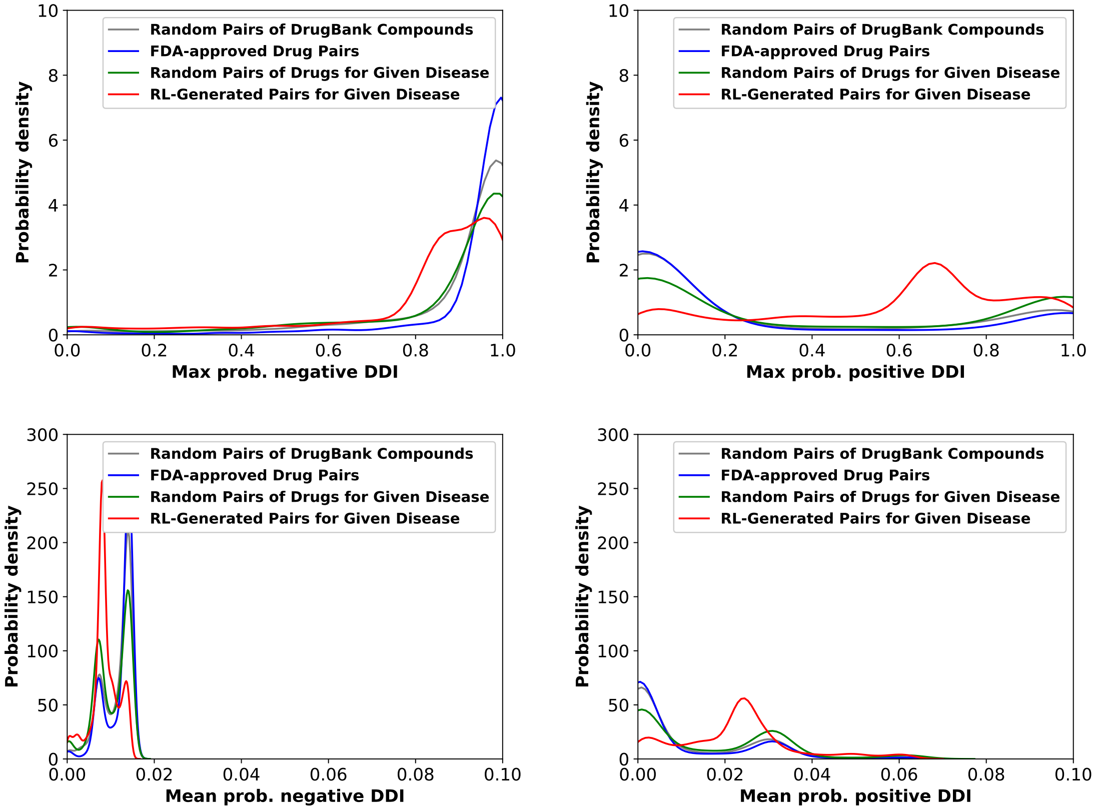


Figure S5: Comparison of distribution of toxicity between RL generated drug combinations, FDA approved drug combinations, randomly selected drug pairs from drugbank and randomly selected drug pairs from FDA approved single drug for ovarian cancer.

##### 3.5 KS Test statistics

###### 3.5.1 Max prob. negative DDI distributions

Table S4: One-sided KS test statistics for comparison of toxicity distributions (p-values)

| RL vs. | drugbank | FDA drug comb. | cancer spec. |
|---|---|---|---|
| Melanoma | 1.01 e-56 | 8.16 e-147 | 4.89 e-92 |
| Breast cancer | 3.82 e-55 | 2.33 e-139 | 1.16 e-91 |
| Lung cancer | 1.88 e-40 | 8.63 e-122 | 5.21 e-74 |
| Ovarian cancer | 2.40 e-98 | 3.75 e-191 | 1.90 e-94 |

###### 3.5.2 Mean prob. negative DDI distributions

Table S5: One-sided KS test statistics for comparison of toxicity distributions (p-values)

| RL vs. | drugbank | FDA drug comb. | cancer spec. |
|---|---|---|---|
| Melanoma | 1.65 e-166 | 2.11 e-162 | 4.02 e-133 |
| Breast cancer | 7.86 e-59 | 1.85 e-86 | 1.81 e-53 |
| Lung cancer | 6.46 e-79 | 6.83 e-101 | 2.08 e-91 |
| Ovarian cancer | 5.97 e-133 | 3.31 e-131 | 1.20 e-66 |

###### 3.5.3 Max prob. positive DDI distributions

Table S6: One-sided KS test statistics for comparison of toxicity distributions (p-values)

| RL vs. | drugbank | FDA drug comb. | cancer spec. |
|---|---|---|---|
| Melanoma | 2.08 e-184 | 9.36 e-173 | 3.66 e-143 |
| Breast cancer | 2.08 e-99 | 1.41 e-124 | 1.52 e-88 |
| Lung cancer | 5.85 e-131 | 2.48 e-137 | 1.17 e-129 |
| Ovarian cancer | 9.57 e-175 | 4.38 e-160 | 4.93 e-90 |

###### 3.5.4 Mean prob. positive DDI distributions

Table S7: One-sided KS test statistics for comparison of toxicity distributions (p-values)

| RL vs. | drugbank | FDA drug comb. | cancer spec. |
|---|---|---|---|
| Melanoma | 2.67 e-197 | 5.70 e-185 | 1.07 e-155 |
| Breast cancer | 8.87 e-102 | 3.79 e-130 | 8.55 e-93 |
| Lung cancer | 6.01 e-128 | 1.18 e-136 | 3.99 e-127 |
| Ovarian cancer | 6.48 e-170 | 7.40 e-161 | 2.06 e-89 |
